# Shallow evolutionary divergence between two Andean hummingbirds: Speciation with gene flow?

**DOI:** 10.1101/249755

**Authors:** Catalina Palacios, Silvana García-R, Juan Luis Parra, Andrés M. Cuervo, F. Gary Stiles, John E. McCormack, Carlos Daniel Cadena

## Abstract

Ecological speciation can proceed despite genetic interchange when selection counteracts homogeneizing effects of migration. We tested predictions of this divergence-with-gene-flow model in *Coeligena helianthea* and *C. bonapartei*, two parapatric Andean hummigbirds with marked plumage divergence. We sequenced neutral markers (mtDNA and nuclear ultra conserved elements) to examine genetic structure and gene flow, and a candidate gene (*MC1R*) to assess its role underlying divergence in coloration. We also tested the prediction of Glogers’ rule that darker forms occur in more humid environments, and compared ecomorphological variables to assess adaptive mechanisms potentially promoting divergence. Genetic differentiation between species was very low and coalescent estimates of migration were consistent with divergence with gene flow. *MC1R* variation was unrelated to phenotypic differences. Species did not differ in macroclimatic niches but were distinct in ecomorphology. Although we reject adaptation to variation in humidity as the cause of divergence, we hypothesize that speciation likely occurred in the face of gene flow, driven by other ecological pressures or by sexual selection. Marked phenotypic divergence with no neutral genetic differentiation is remarkable for Neotropical birds, and makes *C. helianthea* and *C. bonapartei* an appropriate system in which to search for the genetic basis of species differences employing genomics.

## Introduction

New species often arise when geographic isolation of populations allows for divergence via genetic drift or selection (Mayr 1963; Coyne and Orr 2004). Central to this speciation model are the ideas that geographic isolation restricts gene flow, thus allowing for differentiation, and that speciation without geographic isolation is unlikely because gene flow homogenizes populations (Coyne and Orr 2004). Alternatively, the divergence-with-gene-flow model proposes that speciation is possible without geographic isolation if selection is sufficiently strong to counteract the homogenizing effect of gene flow (Gavrilets 1999; Nosil 2008; Pinho and Hey 2010; Martin et al. 2013; Morales et al. 2017). Under this model, phenotypic differentiation may develop in the face of gene flow owing to divergent selection acting on traits directly associated with reproduction or on traits associated with those involved in reproduction through pleiotropic effects (Schluter 2001; Servedio 2016). Assortative mating or selection against hybrids may further facilitate the completion of reproductive isolation (Coyne and Orr 2004; Fitzpatrick et al. 2009; Schluter 2009).

Several studies provide evidence that natural selection can generate phenotypic divergence among populations despite gene flow (e.g. Smith 1997; Morgans et al. 2014; Fitzpatrick et al. 2015) and this could lead to speciation (Hey 2006; Nosil 2008). However, documenting speciation with gene flow is complicated because of the difficulty of determining whether shared genetic variation between species is a consequence of divergence in the presence of migration or rather an indication of post-speciation hybridization or incomplete linage sorting of gene lineages due to recent divergence (Hey 2006; Pinho and Hey 2010). This difficulty has been partly overcome thanks to the development of coalescent-based tools to estimate migration since divergence between pairs of populations (Hey and Nielsen 2004, 2007; Beerli 2006; Kuhner 2006; Durand et al. 2011). Some studies using such tools have found incomplete lineage sorting as the cause for lack of genetic differentiation (Nosil et al. 2009; Wall et al. 2013; Suh et al. 2015), whereas others support population divergence despite gene flow (Green et al. 2010; Rheindt et al. 2014; Supple et al. 2015; Kumar et al. 2017). However, compelling evidence that population divergence has scaled up to the formation of different species in the face of gene flow remains limited. Nonetheless, the finding that the evolutionary histories of various organisms are characterized by substantial cross-species genetic exchange (e.g. Novikova et al. 2016; Zhang et al. 2016; Kumar et al. 2017) implies that attention should be devoted to understanding the selective mechanisms maintaining species as distinct entities in the face of gene flow.

In birds, plumage traits are often targets of natural selection. This results in adaptations for foraging and flight efficiency (Zink and Remsen 1986), camouflage (Zink and Remsen 1986) or conspicuousness (Endler 1993), thermoregulation (Walsberg 1983), and protection against pathogens (Burtt and Ichida 2004; Goldstein et al. 2004; Shawkey et al. 2007), among others. Because plumage traits are also critical in mate selection and species recognition, plumage divergence may drive lineage diversification (Price 2008; Servedio et al. 2011; Hugall and Stuart-Fox 2012; Maia et al. 2013). A frequently observed pattern in presumably adaptive plumage variation is Gloger’s rule, which states that birds with darker plumage coloration occur in more humid environments than lighter-colored conspecifics (Burtt and Ichida 2004). This pattern is often attributed to adaptation to reduce bacterial degradation of plumage in humid conditions where bacteria are most abundant, because melanin (the pigment responsible for black plumage color) confers resistance against these microbes (Goldstein et al. 2004; Peele et al. 2009; Amar et al. 2014). Because differences in melanic pigmentation can serve as cues for mate choice and species recognition (Uy et al. 2009), adaptive differentiation in plumage coloration might thus drive the origin of reproductive isolation. However, we are unaware of studies explicitly relating the evolution of melanic plumage coloration by natural selection to population divergence or speciation in the presence of gene flow (but see Rosenblum et al. 2017; Pfeifer et al. 2018 for examples of involving skin pigmentation in other animals).

Here, we test the divergence-with-gene-flow model of speciation as an explanation for the evolution of two Andean hummingbird species, *Coeligena helianthea* (Blue-throated Starfrontlet) and *Coeligena bonapartei* (Golden-bellied Starfrontlet). We studied these species because: (1) they have largely parapatric ranges in a topographically complex area of the Andes over which environmental conditions (hence selective pressures) may differ (Fig. 1). (2) They lack genetic differentiation in neutral markers (Parra et al. 2009; McGuire et al. 2014) as expected under divergence with gene flow. (3) They exhibit distinct phenotypic differences (plumage in *C. helianthea* is considerably darker than in *C. bonapartei*) and no hybrids have been reported even where they coexist locally (except perhaps for a few old specimens; Fjeldså & Krabbe, 1990). And (4), because variation in melanic pigmentation may reflect adaptation to different environments, differentiation in plumage traits between these hummingbird species might have been driven by natural selection.

**Figure 1.**
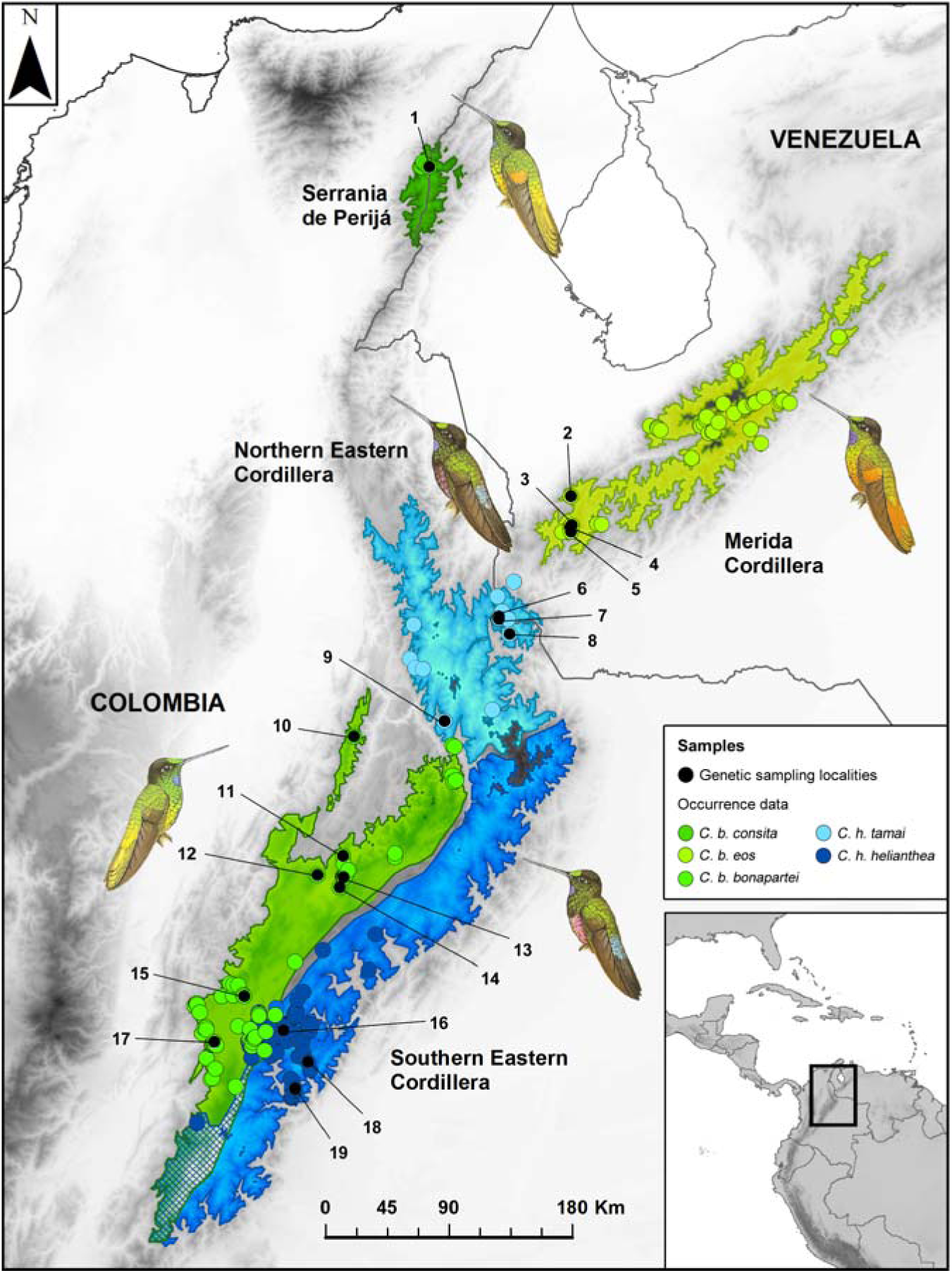
Geographical distribution and sampled localities of *C. helianthea* and *C. bonapartei*. Black dots correspond to localities of specimens sampled for genetic markers. Colored dots correspond to occurrence data obtained from public data bases (see Material and Methods). Both were used for niche overlap analysis. Polygons correspond to the likely distributions of the subspecies according to elevational limits (Ayerbe-Quiñones 2015) and occurrence data.

The apparent lack of genetic differentiation between *C. helianthea* and *C. bonapartei* (Parra et al. 2009; McGuire et al. 2014) despite their distinct differences in potentially adaptive traits may reflect divergence with gene flow, contemporary hybridization, or incomplete lineage sorting (Hey 2006; Suh et al. 2015; Sonsthagen et al. 2016). We here evaluate predictions of the divergence-with-gene-flow model of speciation and consider the evolutionary mechanisms driving divergence between these species by first addressing the following questions: (1) does the lack of genetic differentiation between *C. helianthea* and *C. bonapartei* persist with a much larger and geographically extensive sampling and additional molecular markers relative to earlier work (Parra et al. 2009)?, and (2) are patterns of genetic variation consistent with a model of divergence in the face of gene flow? We next asked (3) is color divergence associated with genetic variation in the *MC1R* gene, a candidate underlying melanic coloration in various bird species and other vertebrates? To examine possible mechanisms through which natural selection might have driven population differentiation we examined whether phenotypic divergence may be attributable to adaptation to contrasting macro-environmental conditions by asking (4) is *C. helianthea* with darker plumage distributed in more humid environments as predicted by Gloger’s rule? and (5) is there morphometric variation between species that may suggest adaptations to alternative microhabitats or resources?

## Materials and Methods

### Study system

*Coeligena helianthea* inhabits mostly the eastern slope of the Cordillera Oriental of the Northern Andes from western Meta in Colombia to the Táchira Depression in Venezuela, and comprises two subspecies: *C. h. helianthea* occupies most of the range, whereas *C. h. tamai* occurs in the Tamá Massif in the border between Colombia and Venezuela (Fig. 1). The distribution of *Coeligena bonapartei* is not continuous and three subspecies are recognized: 1) *C. b. bonapartei* ranges along the western slope of the Cordillera Oriental in Cundinamarca, Boyacá, and western Santander in Colombia, 2) *C. b. consita* is restricted to the Serranía del Perijá, and 3) *C. b. eos* is endemic to the Cordillera de Mérida in the Venezuelan Andes (Hilty and Brown 1986; Hilty 2003. Fig. 1). Some authors consider the Venezuelan taxon *C. b. eos* a distinct species (Del-Hoyo et al. 1999; Donegan et al. 2015), but it is currently treated as a subspecies of *C. bonapartei* (Remsen et al. 2017). Although the distributions of *C. bonapartei* and *C. helianthea* are not sympatric for the most part, the nominate subspecies co-occur regionally in Cundinamarca and Boyacá (Gutiérrez-Zamora 2008).

*C. helianthea* and *C. bonapartei* differ strikingly in plumage coloration. Although both species have bright green crowns and violet gorgets, males of *C. helianthea* are considerably darker, with a largely greenish black with a rose belly and aquamarine rump; males of *C. bonapartei* are largely golden green with fiery gold underparts and rump. Females are paler than males, but also differ distinctly in plumage, especially in their lower underparts (Hilty and Brown 1986; Parra 2010). The differences in coloration between species may reflect variation in the melanin content of feathers (D’Alba et al. 2014), but may also be due to differences in the nanostructure of feather barbules (i.e. width of the air spaces or keratin layer), which interferes with light to generate the reflected colors (Greenewalt et al. 1960).

### Tissue samples and DNA sequencing protocols

We collected specimens in Colombia and Venezuela, and obtained tissue samples from the collections of the Instituto Alexander von Humboldt (IAvH), the Museo de Historia Natural de la Universidad de los Andes (ANDES), and the Colección Ornitológica Phelps (Table S1). Our sampling included a total of 62 individuals: 38 specimens of *C. bonapartei* (12 *C. b. bonapartei*, 5 *C. b. consita*, and 21 *C. b. eos*) and 24 specimens of *C. helianthea* (7 *C. h. helianthea*, 17 *C. h. tamai*). Subspecies were assigned based on taxonomic determination of museum specimens or by geography. We extracted DNA from tissue samples using either a QIAGEN DNeasy Tissue Kit (Qiagen, Valencia, CA, USA) following the manufacturer’s instructions or a standard phenol/chloroform extraction protocol. For 60 specimens we amplified by PCR (Methods S1) and sequenced 1041 bp of the mitochondrial *ND2* gene, and used the data for range-wide phylogeographic and population genetic analyses. We used published sequences of *C. lutetiae* (McGuire et al. 2007; Parra et al. 2009) and *C. orina* (McGuire et al. 2014) as outgroups in phylogenetic analyses.

We used a subset of 36 individuals to assess whether color differentiation between species is associated with nucleotide substitutions in the coding region of the *melanocortin-1 receptor* gene (*MC1R*), a locus responsible for melanic pigmentation in several birds and other vertebrates (Mundy 2005; Roulin and Ducrest 2013). We amplified by PCR (Methods S1) and sequenced 788 bp of the 945 bp of the *MC1R* locus for 6 individuals of *C. h. helianthea*, 10 *C. h. tamai*, 8 *C. b. bonapartei*, 1 *C. b. consita* and 11 *C. b. eos*. All PCR products were cleaned and sequenced in both directions by Macrogen Inc. or at the sequencing facilities of the Universidad de los Andes. We assembled, edited, and aligned sequences of the *ND2* and *MC1R* genes using BioEdit 7.2.5 (Hall 1999) and Geneious 9.1.5 (http://www.geneious.com/; Kearse et al., 2012), employing the MUSCLE algorithm and manual editing.

We also employed a sequence capture approach to acquire data from regions flanking ultraconserved elements (UCEs; Faircloth et al. 2012) for 1 individual of *C. h. helianthea*, 4 *C. h. tamai*, 1 *C. b. bonapartei*, and 1 *C. b. consita* to obtain a preliminary overview of genetic divergence between these taxa at a genomic level. We used a standard library preparation protocol (http://ultraconserved.org/; Faircloth & Glenn, 2012) and enriched the pool of samples for 5,060 UCE loci using the MYbaits_Tetrapods-UCE-5K probes. We sequenced the pool after quantification using Illumina MiSeq. Following the PHYLUCE pipelines (Faircloth 2015), we used Illuminoprocessor (Faircloth 2013) and Trimmomatic (Bolger et al. 2014) to trim reads, discarded adapter contamination and low-quality bases, and assembled the reads into contigs using a kmer=50 and ABySS (Simpson et al. 2009). We aligned the contigs against the original UCE probes to identify contigs matching UCE loci using LASTZ (Harris 2007). Among the individuals, we aligned UCE loci using the default MAFFT v7.13 algorithm (Katoh and Standley 2013). Finally, we pulled out UCE loci from the Anna’s Hummingbird (*Calypte anna*) genome (Gilbert et al. 2014; Zhang et al. 2014) to use them as outgroup.

For phylogenetic analysis we used a concatenated alignment of 2,313 UCE loci shared at least among 3 individuals including the outgroup. Of these, 1,465 loci were present in all the individuals (mean locus length = 615.1 bp, mean number of individuals per locus in the incomplete matrix = 7.3). We generated a second concatenated alignment of 1,604 loci shared among all *Coeligena* specimens (i.e. without the outgroup). Of these, 389 loci showed no variation, 75 had only indels (informative or not), 614 had singletons and indels, and 526 (32.8%) had informative sites (polymorphic sites with each variant represented in at least two individuals). We used the latter 526 loci for population genetic analyses aimed at assessing gene flow.

### Phylogenetic and Population Genetic Analysis

We used maximum-likelihood and Bayesian inference methods to reconstruct phylogenies from the CIPRES Portal (http://www.phylo.org/) or locally. We conducted maximum-likelihood analyses in RAxML (Stamatakis 2014) using the GTR+GAMMA model and non-parametric bootstrapping under the autoMRE stopping criterion for *ND2* and UCE data. We conducted Bayesian analyses in Mr. Bayes v3.2 (Ronquist et al. 2012) using a single partition and the HKY model for *ND2* data, which was the best fit according to JModelTest 2.1.7 (Posada 2008; Darriba et al. 2012). For UCE data we used 16 partitions and the models for each of these suggested by CloudForest analysis (Crawford and Faircloth 2011). The MCMC parameters consisted of two runs with four chains ran for 15 million generations sampling every 100 generations for the *ND2* data, and ran for 25 million generations sampling every 500 generations for the UCE data. We discarded the first 10% generations as burn-in before estimating the consensus tree and posterior probabilities. Convergence and effective sample sizes of parameter estimates were examined using Tracer 1.6 (Rambaut et al. 2016).

To further examine relationships among *ND2* haplotypes, we used an alignment of 885 bp for which complete data were available for all individuals to construct a haplotype network in Network 5.0.0.1 (http://www.fluxus-engineering.com/; Bandelt, Forster, & Röhl, 1999). To examine genetic structure between species, we calculated Fst with R package hierfstat (Goudet and Jombart 2015) and AMOVAs with R package ade4 (Dray and Dufour 2007) assessing significance using 10,000 permutations (Script S1). Also, we used the program Structure 2.3.4 (Pritchard et al. 2000) to assess population structure using UCE data. We performed 20 runs for each value of K from K=1 to K=5, using a burnin period of 10,000 steps and 100,000 repetitions.

We followed Pritchard et al. (2000) to calculate the probability of different values of K using the mean ln likelihood value calculated over the 20 runs as prob(K=n) = (elnK=n) / (elnK=1+…+elnK=n).

### Testing for Divergence with Gene Flow

We used Migrate 3.2.1 (Beerli 2009) to examine whether lack of genetic differentiation observed between *C. helianthea* and *C. bonapartei* is more likely a result of speciation in the face of gene flow or rather a consequence of hybridization following secondary contact. In addition to estimating parameters such as effective population size scaled by mutation rate (θ) and migration scaled by mutation rate (M), Migrate can estimate parameters for different time bins, allowing one to estimate migration at different moments through time. If there has been gene flow between species after speciation, then posterior distributions of migration estimated as M=(m/µ) should exclude values of zero. Given non-zero estimates of migration, divergence-with-gene-flow predicts higher values of M close to the time of divergence, whereas post-speciational gene flow (i.e. recent hibridization) predicts higher values of M near the present.

We used two data sets for Migrate analyses. First, we employed a *ND2* alignment of 885 bp (excluding all the positions with missing data) including 22 individuals of *C. helianthea* and 17 individuals of *C. b. bonapartei/consita* (i.e. excluding *C. b. eos*, which we found to be genetically distinct; see below). Second, we used an alignment of 591 SNPs (368 informative sites) derived from 296 UCE loci; we used only those UCE loci having data for all seven individuals and considered only sites where SNPs showed variation between at least two individuals. Because inference of gene flow requires using markers that have evolved neutrally, we first confirmed that both data sets meet this assumption by calculating Tajima’s D using DNAsp 5.1 (Librado and Rozas 2009).

We determined prior maximum values for the parameters θ and M for each species and each data set based on several test runs. In final analyses aimed to estimate gene flow using both *ND2* and UCE data, we set prior values to 0.1 for θ for both species, and to 1,000 for M from *C. bonapartei* to *C. helianthea* and to 600 for M from *C. helianthea* to *C. bonapartei*. We ran Migrate in the CIPRES Portal (http://www.phylo.org/) using a long chain of 300 million steps (sampling 100,000 steps recorded every 3,000 steps) with a burn-in of 100,000 steps.

### *MC1R* gene analyses

We compared variable sites in *MC1R* sequences between our study species and translated sequences to aminoacids to check for synonymous and non-synonymous substitutions. As references for comparisons we used sequences of *Calypte anna* and Chimney Swift (*Chaetura pelagica*) predicted from genome annotations (Zhang et al. 2014). Because these comparisons revealed no variation potentially implied in phenotypic variation (see results), we did not conduct any additional analysis.

### Examining the selective regime: niches and ecomorphological differentiation

We tested the hypothesis that natural selection underlies the phenotypic divergence in color between *C. heliathea* and *C. bonapartei* through macroclimatic differences in the regions occupied by these species. Specifically, we tested the prediction of Gloger’s rule that *C. helianthea* (with darker plumage) occurs in environments with more humid conditions than *C. bonapartei*, and examined whether other macroclimatic conditions that may promote adaptation differ between environments occupied by these hummingbirds. We examined ecological differentiation among *C. helianthea*, *C. b. bonapartei/consita* and *C. b. eos* (which we found to be genetically distinct; see below) using occurrence data, environmental variables, and measurements of niche overlap (Broennimann et al. 2012). In addition to the locality data associated with specimens included in molecular analyses, we obtained occurrence data from eBird (http://ebird.org/content/ebird/), Vertnet (http://vertnet.org/), GBIF (http://www.gbif.org/), Xeno-canto (http://www.xeno-canto.org/), and the ornithological collection of the Instituto de Ciencias Naturales of the Universidad Nacional de Colombia (http://www.biovirtual.unal.edu.co/en/), for a total of 242 records. After eliminating duplicates and excluding non-reliable locations we retained 196 records for analysis: 85 of *C. helianthea*, 75 of *C. b. bonapartei/consita*, and 36 of *C. b. eos*.

To delimit the accessible areas for each species we used ecoregions as defined by Dinerstein et al. (Dinerstein et al. 2017). We used all the ecoregions with occurrence records as the environmental background available for the analysis of niche overlap. We obtained climatic data from WorldClim (http://www.worldclim.org/ Hijmans et al. 2005), CliMond (https://www.climond.org/ Kriticos et al., 2012), and EarthEnv (http://www.earthenv.org/cloud Wilson and Jetz 2016). We selected and excluded variables highly correlated to others (Threshold: 0.7) using the package usdm (Naimi 2015) in R (R Core Team 2016). We conducted niche overlap analyses using 11 variables: three related to temperature, three related to precipitation, four related to cloudiness, and one related to air moisture (Table S2).

We extracted climatic data from 10,000 points from the background environment and from the 196 occurrence records and performed a principal component analysis (PCA) to summarize climatic variation using the R package ade4 (Dray and Dufour 2007). With the two first PCA axes, we plotted the densities of each taxon in climatic space relative to the background using R package ecospat (Broennimann et al. 2016). We also used this package to estimate the D statistic (Warren et al. 2008) to quantify niche overlap (D = 0 indicates different niches, and D = 1 indicates identical niches), and we performed similarity tests (1,000 iterations) to assess whether niches are less similar (niche divergence) than expected by chance given background climatic variation (Script S2). Significant niche divergence with the darker *C. helianthea* occupying more humid areas would be consistent with adaptive divergence following Gloger’s rule, whereas no significant differences in niches would suggest that adaptation to distinct climatic conditions cannot account for phenotypic differentiation between species.

We also assessed whether there is morphometric differentiation between species which may reflect adaptation to different microhabitats or food resources (Stiles 2008) by measuring 17 traits related to beak, wing, tail and leg morphology (Table S3). We measured morphological variables from 35 live individuals (17 females and 18 males) of *C. h. helianthea* and 46 individuals (23 females and 23 males) of *C. b. bonapartei*. Using these data we asked whether individuals of different species and sexes are distinguishable in multivariate space employing linear discriminant analysis (LDA) using R package MASS (Venables and Ripley 2002). We also built ANOVA models to test for mean differences in individual variables among species and sexes simultaneously (Script S3).

## Results

### Does lack of genetic differentiation between *C. helianthea* and *C. bonapartei* persist with greater sampling and additional markers?

We found low genetic differentiation between *C. b. bonapartei* and *C. b. consita*, but both taxa were markedly differentiated from *C. b. eos*. Therefore, in the following we treat *C. b. bonapartei* and *C. b. consita* as a single group, which we refer to as *C. b. bonapartei/consita*. Divergence in *ND2* of *C. helianthea* and *C. b. bonapartei/consita* relative to *C. b. eos* was high, with siginificant Fst values of 0.56 and 0.52, respectively (p < 0.001 in both cases), and relatively high fractions of genetic variance (59.4% and 52.0%, respectively) existing between groups in AMOVA. In contrast, *ND2* data showed little to no differentiation between *C. helianthea* and *C. b. bonapartei/consita.* Although differentiation as measured by Fst was significant (p = 0.03), the Fst value was very low (0.07) and only 1.7% of the variance was partitioned between these two taxa in AMOVA, with 98.3% of the variance existing among individuals within taxa.

Phylogenetic analyses showed that *C. b. eos* forms a strongly supported clade (posterior probability PP = 1.0, maximum-likelihood bootstrap MLbs = 87%), which is sister to a moderately supported clade (PP = 0.77, MLbs = 69%) formed by *C. helianthea* and *C. b. bonapartei/consita* (Fig. 2A). Within the latter clade, relationships among populations appeared to be determined more by geography than by current species-level taxonomy: most sequences of the northern subspecies *C. b. consita* and *C. h. tamai* formed a strongly supported clade (PP = 1.0, MLbs = 85%), whereas the majority of sequences of southern subspecies *C. b. bonapartei* and *C. h. helianthea* formed another moderately supported clade (PP = 0.94, MLbs = 61%).

**Figure 2.**
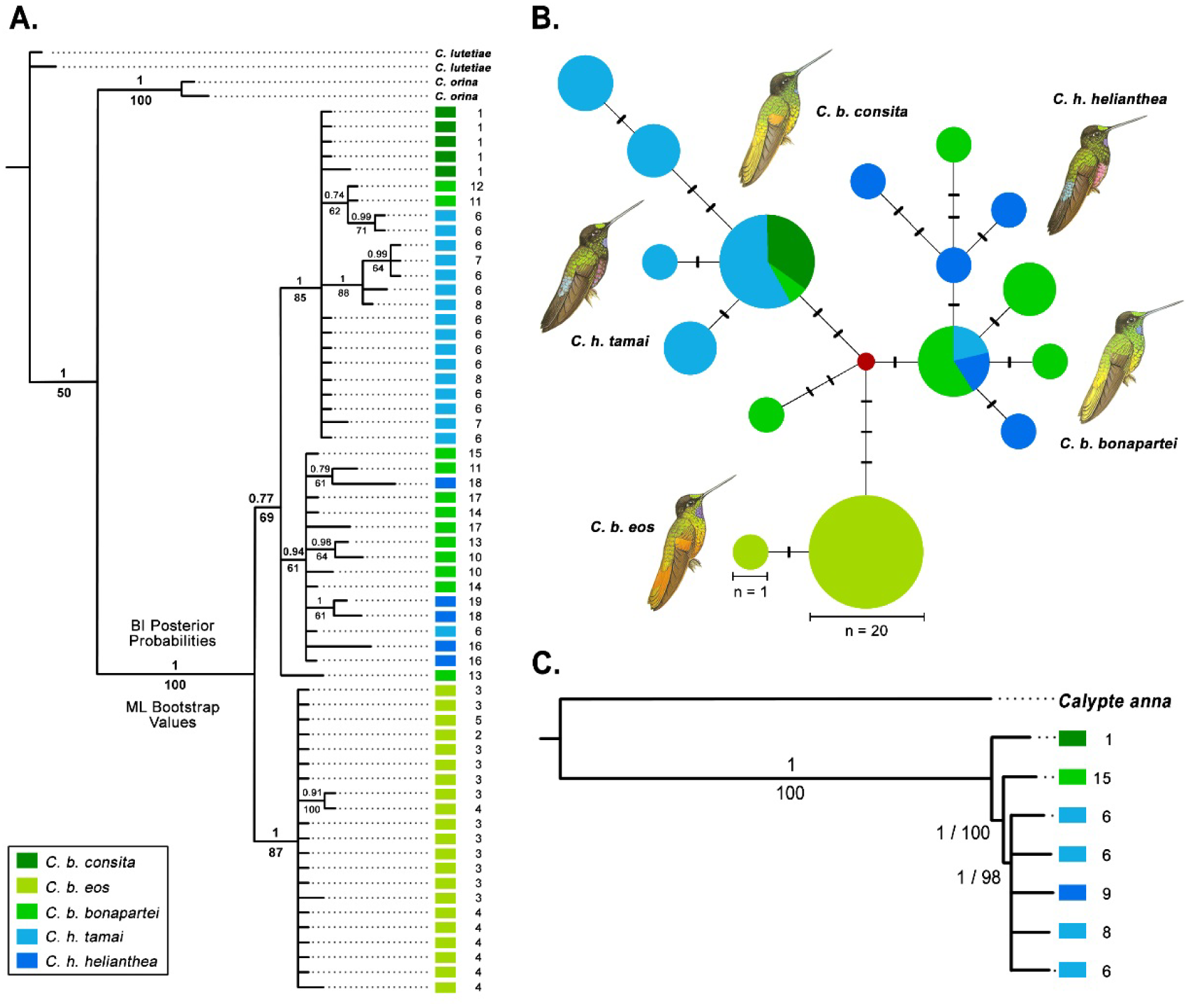
*ND2* phylogenetic reconstructions and haplotype network show lack of divergence between *C. helianthea* and *C. b. bonapartei/consita*. The *ND2* gene trees (A) and haplotype networks (B) show *C. helianthea* and *C. b. bonapartei/consita* in a single clade separate from a *C. b. eos* clade. Most specimens of the northern subspecies *C. h. tamai* and *C. b. consita* cluster together, whereas southern subspecies *C. h. helianthea* and *C. b. bonapartei* form another cluster, suggesting that population structure more strongly reflects geography (i.e. north-south differentiation) than taxonomy based on plumage phenotype. The phylogenetic reconstruction based on UCE loci shows *C. helianthea* nested within *C. b. bonapartei/consita* (C). Numbers at the right of the individuals in the tips of the trees correspond to the sampled localities (Table S1).

Haplotype networks confirmed the above findings (Fig. 2B): (1) *C. helianthea* and *C. b. bonapartei/consita* shared haplotypes, whereas *C. b. eos* did not share any haplotypes with the other taxa; and (2) haplotype groups were more consistent with geography than with taxonomy. However, networks showed that the latter pattern is not perfect because two individuals of *C. b. bonapartei* (from the south) had the haplotype most common in the north, one *C. h. tamai* (from the north) had the haplotype most common in the south, and one *C. b. bonapartei* had an intermediate haplotype.

UCE nuclear markers also did not reveal genetic differentiation between *C. helianthea* and *C. bonapartei*. The UCE phylogeny shows a well-supported clade including all sequences of *C. helianthea* nested within a clade in which the two earliest diverging branches were the two specimens of *C. bonapartei* (Fig. 2C). Aditionally, population genetic structure between species in UCE markers was not significant (Fst = 0.2, p = 0.5), and the most likely number of genetic clusters in the data set according to Structure was K = 1 (prob_(K=1)_ = 0.8). Support for larger values of K was much lower and clusters defined assuming different values of K never corresponded to groups defined by species identity (Fig. S1).

### Are patterns of genetic variation consistent with divergence in the face of gene flow?

Our data sets fit the assumption of neutrality, allowing one to use them for gene flow inference: Tajima’s D values were not significant for either marker (*ND2* D=-0.68 p >0.1, UCEs D=0.22 >0.1). The analysis of both *ND2* and UCE data suggested that there has been gene flow between *C. helianthea* and *C. bonapartei* after their divergence. Mean estimates of migration (M=m/µ) were in all cases different from zero: M = 725.1 from *C. helianthea* to *C. bonapartei* and 446.1 from *C. bonapartei* to *C. helianthea* for *ND2*, and M = 869.9 from *C. helianthea* to *C. bonapartei* and 555.4 from *C. bonapartei* to *C. helianthea* for UCE data. However, the posterior probability distributions of M estimated from the *ND2* data were wide: 95% credibility intervals ranged from 284.7 to 1,000 from *C. helianthea* to *C. bonapartei,* and from 0.0 to 628 from *C. helianthea* to *C. bonapartei* (Fig. 3A). In contrast, posterior distributions of M estimated from UCE data were narrowly concentrated around the mean: 95% credibility intervals ranged from 782.0 to 958.7 from *C. helianthea* to *C. bonapartei* and from 482.0 to 626.7 from *C. bonapartei* to *C. helianthea* (Fig. 3C), rejecting scenarios of no migration after divergence.

**Figure 3.**
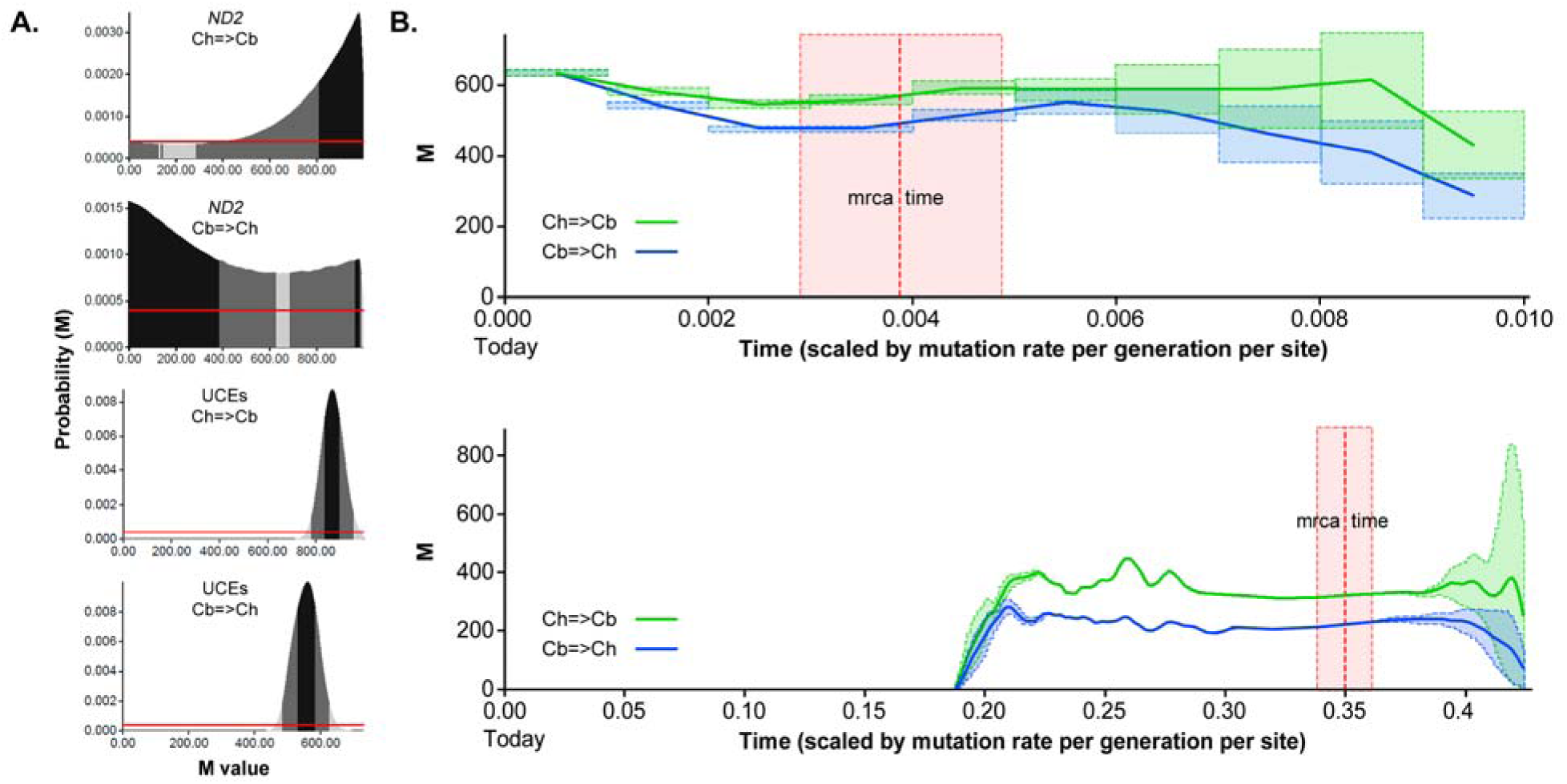
Migration parameter estimates suggest gene flow after the divergence between *C. helianthea* to *C. bonapartei*. Posterior distributions of the migration parameter M=m/µ from *C. helianthea* to *C. bonapartei* and vice versa (A) estimated based on *ND2* (top) and UCE (bottom) data; colors correspond to the limits of the intervals accumulating 50% (black), 75% (dark gray) and 95% (light gray) of the probability density. The red horizontal line corresponds to the prior, which is constant. Migration parameter M estimated value (y axis) from *C. helianthea* to *C. bonapartei* (green) and vice versa (blue) changing through time (scaled by mutation rate per generation per site, 0 = today) (B) for the *ND2* (upper panel) and UCE (bottom panel) data sets. Dashed boxes in green and blue depict ca. 1.96 of standard error of the estimated value of M. The red vertical dashed lines and boxes correspond to the mean value and one standard deviation, respectively, of the estimated time of the most recent common ancestor of the species.

Estimates of migration through time further supported that divergence occurred and has been maintained in the face of gene flow as predicted by the divergence-with-gene-flow model of speciation. Our analyses indicated that migration between *C. helianthea* and *C. bonapartei* continued after their initial divergence (i.e. the estimated time of their most common recent ancestor, Fig. 3B,D). Whereas *ND2* data suggested that gene flow has continued until the present (Fig. 3B), the UCE data suggested that gene flow likely ceased at approximately half the time passed since these species last shared a common ancestor (Fig. 3D).

### Is color divergence associated with genetic variation in *MC1R?*

Of the 36 *Coeligena* individuals sampled for *MC1R*, 32 shared a haplotype (excluding ambiguous positions). Genetic variation at *MC1R* was limited to three individuals of *C. helianthea* and one individual of *C. bonapartei*, and involved changes in four sites. Only one change was non-synonymous (Ser275 [AGC] → Arg275 [AGG] at nucleotide site 825), but it was present in a single *C. helianthea* (Andes-BT 1126) with typical plumage coloration. These results reveal no association between *MC1R* genotype and species-specific color phenotypes in *C. helianthea* and *C. bonapartei*.

### Is *C. helianthea* with darker plumage distributed in more humid environments as predicted by Gloger’s rule?

We found no support for the prediction that the more darkly colored *C. helianthea* occurs in more humid environments than *C. b. bonapartei/consita*: the climatic niches of these taxa overlap considerably (D = 0.65, Fig. 4A) and we found no evidence for significant niche divergence relative to background climate (p = 0.99). Niche overlap between *C. b. eos* and *C. b. bonapartei/consita* and *C. helianthea* was considerably lower (D = 0.07 and 0.10, respectively, Fig. 4B), but relative to the background niche differences were not significant (p = 0.70 and 0.76, respectively).

**Figure 4.**
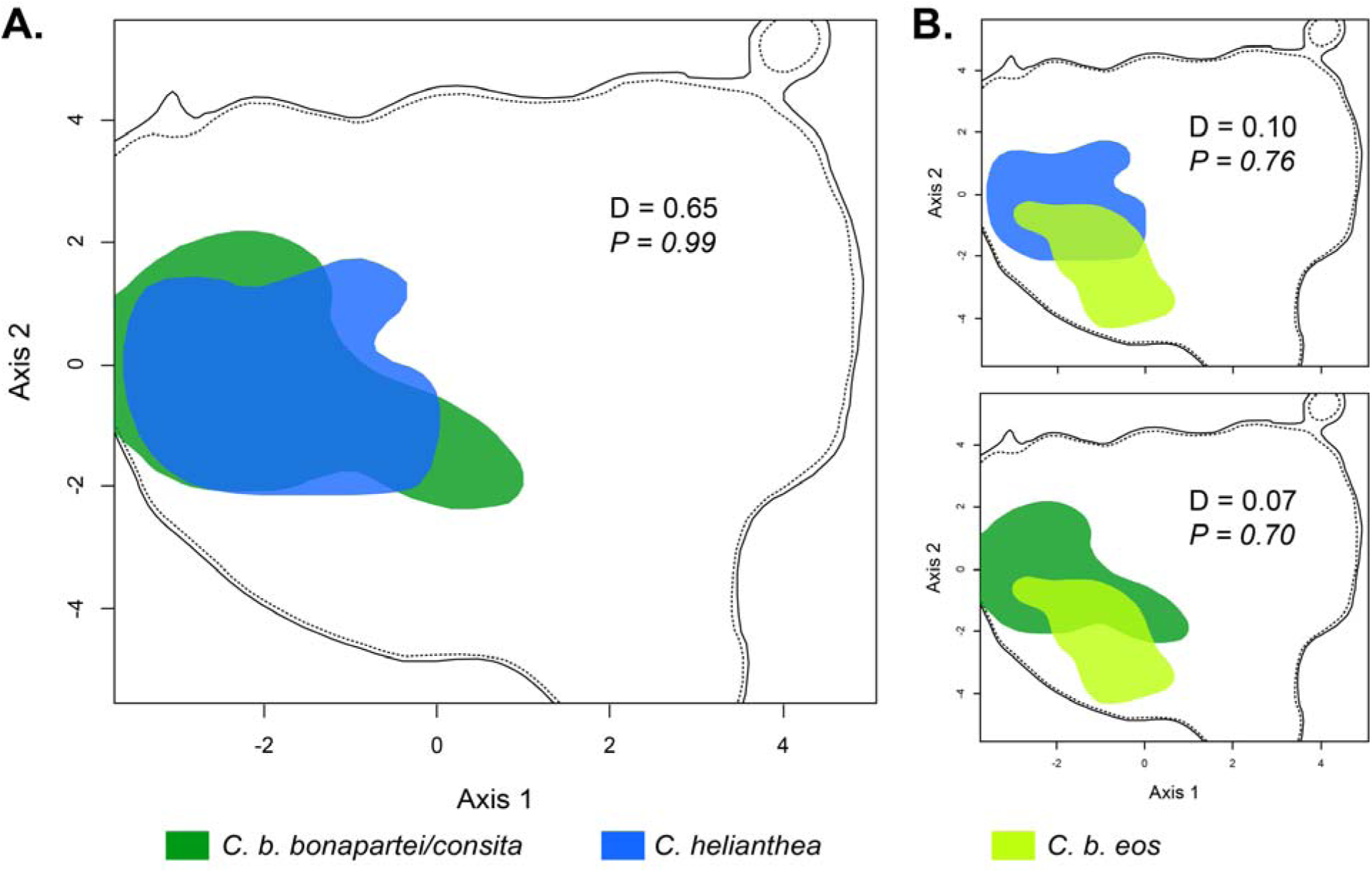
*C. helianthea* and *C. b. bonapartei/consita* do not differ in climatic niches thus do not support Gloger’s rule. The climatic niches of *C. helianthea* and *C. b. bonapartei/consita* overlap considerably (D = 0.65) (A). The climatic niche of *C. b. eos* overlaps very little with *C. helianthea* (D = 0.10) and *C. b. bonapartei/consita* (D = 0.07) climatic niches (B). Nevertheless, relative to the background the differences between the niches are not significant in any case (p>0.1).

### Is there morphometric variation between species that may suggest adaptations to alternative microhabitats or resources?

Morphometric data showed differences between *C. h. helianthea* and *C. b. bonapartei* and between females and males of each taxon: LDA analysis distinguished species/sex with a low classification error of 1.2%. The two most relevant variables in the LD function were wing loading (coefficients: LD1 = 240.2, LD2 = 287.8, and LD3 = 120.0), and wing taper (coefficients: LD1 = −39.7, LD2 = −42.6, and LD3 = −23.4). ANOVA models showed significant differences in 12 morphological variables between species, and in 15 variables between sexes (see Fig. S2). The three variables that differed the most between species and sexes were length of extended wing (ANOVA coefficients: −3.4 species and 5.9 sex), total culmen (ANOVA coefficients: 2.3 species and 2.2 sex), and length of tail (ANOVA coefficients: 1.0 species and 3.7 sex). *Coeligena b. bonapartei* has longer wings, shorter bills and shorter tails than *C. h. helianthea* (p = < 0.001 in all cases), and females have shorter wings, longer bills and shorter tails than males in both species (p = < 0.001 in all cases). Our analyses further revealed that the magnitude of morphometric differences between sexes varied by species. For example, females of *C. helianthea* are the smallest of the four groups (i.e. combinations of species and sexes), but males of *C. helianthea* are the largest.

## Discussion

*C. helianthea* and *C. bonapartei* are sister species of hummingbirds from the Northern Andes that differ distinctly in plumage coloration, but we found a striking lack of genetic differentiation between them in a mitochondrial gene (*ND2*) and in 1,604 UCE markers broadly scattered across the genome. The strong phenotypic differences between *C. helianthea* and *C. bonapartei* in the absence of neutral genetic differentiation are remarkable for Neotropical birds, and make these species an appropriate system in which to search for the genetic basis and adaptive significance of phenotypic differences involved in speciation (see Campagna et al. 2017). However, we found no evidence that *MC1R* (a candidate gene associated with melanic pigmentation in a variety of vertebrates) underlies phenotypic variation, and found no support for the hypothesis that Gloger’s rule (adaptation to geographic variation in humidity) or other macroclimatic niche differences are associated with phenotypic divergence between these species. Nonetheless, coalescent estimates of migration indicate that *C. helianthea* and *C. bonapartei* diverged in the presence of gene flow, suggesting that phenotypic differences likely originated under selective pressures strong enough to offset the homogenizing effects of migration. Our finding that *C. h. helianthea* and *C. b. bonapartei* differ in morphometric traits related to habitat and resource use is consistent with the hypothesis that natural selection may have played a role in their divergence. In addition, as we discuss below, phenotypic divergence may have been maintained in the face of gene flow because of sexual selection.

Although shallow genetic divergence between species may also result from processes including incomplete lineage sorting or contemporary hybridization after secondary contact, our coalescent analyses are consistent with a scenario where *C. helianthea* and *C. b. bonapartei/consita* (i.e. excluding *C. b. eos*) have exchanged genes since their divergence from a common ancestor. Our inference of gene flow is likely robust because both *ND2* and UCE data fit neutrality (Hey and Nielsen 2004). Thus, our data are consistent with the hypothesis that speciation occurred in the face of gene flow, with divergence in loci underlying phenotypic differences between species likely maintained by some form of selection. Future work should conduct analyses examining other markers across the genome because this may allow reducing uncertainty in the estimation of population genetic parameters (Hey and Nielsen 2004), may help rule out other processes such as incomplete lineage sorting (Suh et al. 2015), and may allow one to identify the genetic basis of phenotypic differences (Bourgeois et al., 2016, Toews et al. 2016, Campagna et al. 2017).

We found no variation between species in the coding region of *MC1R*, a gene associated with variation in plumage coloration in several other birds (Theron et al. 2001; Doucet et al. 2004; Mundy 2004; Baião et al. 2007; Gangoso et al. 2011). Thus, as with other studies showing no association between plumage coloration and variation in *MC1R* (MacDougall-Shackleton et al. 2003; Cheviron et al. 2006; Haas et al. 2009), our work suggests that differences in coloration between *C. helianthea* and *C. bonapartei* are controlled by other genes such as agonists or antagonists of *MC1R* in the melanin metabolic pathway, regions regulating the expression of *MC1R* or other genes (Theron et al. 2001), or genes controlling the shape of the keratin medullar matrix of the feather barb’s spongy layer, which determines light scattering to produce structural colors (Shawkey et al. 2014).

We found no support for Gloger’s rule because the darker *C. helianthea* does not occur in more humid environments than the more lightly colored *C. bonapartei*. Nevertheless, adaptation to different environmental conditions may occur at a finer scale, where habitat differences might select for plumage traits that, for instance, stand out from the background augmenting signal efficacy (Endler 1993; Brumfield and Braun 2001). Indeed, we found that the species differ in morphometric traits (e.g. *C. bonapartei* has longer wings and shorter tails than *C. helianthea*) typically associated with use of different microhabitats or foraging behaviors. Variation in such traits can affect flight speed or the relative ability to maneuver in open vs closed environments (Altshuler et al. 2010; Ortega-Jimenez et al. 2014). To the extent that morphological differences may reflect adaptations to different resources between species (Altshuler and Dudley 2002) and between sexes within species of hummingbirds (Temeles and Kress 2010), our data are consistent with a role for selection driving ecomorphological divergence, but the adaptive value of phenotypic variation, if any, remains to be discovered. Considering that *C. bonapartei* often occurs along forest edges whereas *C. helianthea* is more frequently found in forest interior (Hilty and Brown 1986), studies of the functional consequences of phenotypic differences would be especially useful to assess any potential role of natural selection in driving and maintaining divergence.

Knowledge of the timing of speciation might allow one to make inferences about historical processes that could have promoted divergence between *C. helianthea* and *C. bonapartei*. We can place an upper boundary on the divergence time between these species based on their divergence from *C. lutetiae*, which is 1.6% divergent in *ND2* sequences (Parra et al., 2009). Assuming 2% divergence in mtDNA is approximately equivalent to one million years of isolation (Weir and Schluter 2008), *C. helianthea* and *C. bonapartei* likely split within the past ~ 800,000 years. This time period involves some of the last Pleistocene glaciations when high-altitude environments were uninhabitable and forests likely retreated, resulting in the isolation and divergence of populations (Vuilleumier 1969; Ramírez-Barahona and Eguiarte 2013). It thus remains possible that different selective regimes promoted speciation in these hummingbirds if their divergence occurred across environments with contrasting climatic conditions in the Pleistocene even if they occupy similar environments at present. Although such a hypothesis might be partly testable by modeling historical climates and potential distributions, however, one would still be faced with the question of what evolutionary forces might maintain *C. helianthea* and *C. bonapartei* as distinct given that they occur in regional sympatry in the same macroenvironments in the present.

An alternative explanation for the origin and maintenance of phenotypic distinctiveness in plumage, given the strong sexual dichromatism in *C. helianthea* and *C. bonapartei*, is that their differentiation may have proceeded in the face of gene flow due to sexual selection (Price 1998, 2008). Sexual selection is thought to be a powerful force driving speciation in birds and other organisms (Campagna et al. 2012, 2017; Harrison et al. 2015), and some examples exist of speciation due to sexual selection with gene flow (Servedio 2016). Of direct relevance to our system, a study comparing sexually selected (i.e. gorget and crown coloration) and non-sexually selected traits among *Coeligena* species found that sexual selection may be an important driver of phenotypic differentiation, but that it is probably insufficient for speciation to be completed unless it acts in concert with natural selection (Parra 2010; see also Servedio and Boughman 2017). To assess the plausibility of the hypothesis that sexual selection is involved in the divergence and speciation of *C. helianthea* and *C. bonapartei*, one should test for associations among components of males’ fitness, signaling traits (i.e. coloration, songs), and female preferences. Genomic analyses examining whether there are genetic and signatures of selection acting on regions associated with sexual traits (Charlesworth 2009; Huang and Rabosky 2015; Kirkpatrick 2017) would further help to test the hypothesis of divergence driven by sexual selection.

In conclusion, our study provides evidence that the formation of two species of Andean hummingbirds likely occurred in the face of gene flow, suggesting some form of selection played a role maintaining phenotypic differences and driving speciation. However, because the main selective mechanism we examined (i.e. adaptation to contrasting macroclimatic conditions) appears not to operate in *C. helianthea* and *C. bonapartei*, we conclude that ecological pressures that we did not consider directly or sexual selection were likely involved in their divergence. Future studies should thus aim to test predictions of hypotheses of natural and sexual selection acting on this system. Regardless of the selective processes involved, in line with previous research, our study suggests that selection has played an important role in maintaining phenotypic differences that could lead to speciation in tropical montane birds (Cadena et al. 2011; Winger and Bates 2015). Finally, the shallow genetic divergence that we observed between these species suggest that their genomes are unlikely to have been substantially affected by processes occurring after speciation (e.g. post-speciation divergence by drift), which makes this system especially promising for work on the genomics of speciation. Studies aiming to understand the genetic underpinnings of species differences employing genomic approaches (e.g. Campagna et al. 2017; Stryjewski and Sorenson 2017) will be an important complement to our increasing knowledge of the geographic and ecological context of speciation in tropical montane birds.

## Acknowledgements

We thank the Facultad de Ciencias at Universidad de Los Andes for financial support through the Proyecto Semilla program. For providing tissue samples and supporting fieldwork, we thank the Museo de Historia Natural de la Universidad de los Andes (ANDES), the Instituto Alexander von Humboldt (IAvH) and the Colección Ornitológica Phelps (Jorge Perez-Emán and Miguel Lentino). We also thank Whitney Tsai, Paola Montoya, Camila Gomez and Laura Céspedes for their help with laboratory work and analyses. The manuscript was improved thanks to comments by L. Campagna and E. Jarvis, and discussion with Irby Lovette’s and Daniel Cadena’s lab groups.

